# Complete male-to-female sex reversal in XY mice lacking the *miR-17∼92* cluster

**DOI:** 10.1101/2023.03.22.533123

**Authors:** Alicia Hurtado, Irene Mota-Gómez, Miguel Lao, Francisca M. Real, Johanna Jedamzick, Miguel Burgos, Darío G. Lupiáñez, Rafael Jiménez, Francisco J. Barrionuevo

## Abstract

In mammals, sex determination is controlled by antagonistic gene cascades operating in embryonic undifferentiated gonads^1^ ^2^. The expression of the Y-linked gene *SRY* is sufficient to trigger the testicular pathway, whereas its absence in XX embryos leads to ovarian differentiation^3^ ^4^ ^5^. Despite this strong genetic component, the involvement of non-coding regulation in determining mammalian sex remains unclear^6^. Here we show that the deletion of a single microRNA cluster, *miR-17∼92*, induces complete primary male-to-female sex reversal in XY mice. Time-course analyses revealed that *Sry* is heterochronically expressed, showing a delay in XY *miR-17∼92* knockout gonads, which subsequently activate the ovarian genetic program. Bulk and single cell RNA-seq analyses showed that Sertoli cell differentiation is reduced, delayed and unable to sustain the testicular fate. This disrupted differentiation results from a transient state of sex ambiguity in pre-supporting cells, which is later resolved towards the ovarian fate. Consistent with known mechanisms of miRNA-mediated gene regulation, the expression of *miR-17∼92* target genes is not stabilized in undifferentiated XY mutant gonads, affecting concomitantly the fine regulation of gene networks with critical roles in developing gonads. Our results demonstrate that microRNAs are key components for mammalian sex determination, controlling the timing of *Sry* expression and Sertoli cell differentiation.

## INTRODUCTION

Mammalian sex determination involves the simultaneous expression of genes with antagonistic functions^1^ ^2^, resulting in a balanced network of opposing female-and male-promoting signalling factors. The Y-linked gene, *SRY,* is the trigger that breaks this balance: its expression in the undifferentiated gonads from XY individuals induces testis differentiation, whereas its absence in XX gonads results in ovarian development^3^ ^4^ ^5^. In this bipotential system, the repression of genes promoting the alternate fate is paramount for sex-specific differentiation^1^ ^7^ ^8^. Further, this mutual antagonism is considered as a landmark of both mammalian sex determination and maintenance, highlighting the strong genetic component that influences this developmental pathway.

Previous research in the sex determination field has mostly focused on the identification of genes controlling this process and on elucidating their hierarchical relationships at a molecular level. Yet, the involvement of non-coding regulatory mechanisms in mammalian sex determination remain largely unexplored^9^ ^6^. One intriguing class of non-coding regulators are microRNAs (miRNAs), short nucleotide sequences that mediate the post-transcriptional regulation of gene expression. These small molecules were initially characterized as temporal regulators of cell-fate decisions^10^ ^11^ ^12^: their inactivation led to heterochronic shifts in gene expression and differentiation. Progressively, miRNAs have been also shown to be implicated in numerous developmental and pathological processes^13^. During sex development, both miRNAs and their associated machinery are differentially regulated in developing testes and ovaries^14^. Furthermore, the disruption of their biosynthesis in adult mice compromised their fertility, confirming a role of miRNAs in maintaining gonadal cell type identities^15^. However, the inactivation of *Dicer1*, an essential component for miRNA biosynthesis, prior to the sex determination stage did not affect this process^16^. However, the long-time persistence of miRNAs upon disruption of their biosynthesis machinery may have obscured their potential involvement in sex determination^16^.

One of the most well-studied microRNAs is the *miR-17∼92* cluster. This polycistronic cluster, also called OncomiR-1, contains six miRNA genes (*miR-17*, *miR-18a*, *miR-19a*, *miR-20a*, *miR-19b-1* and *miR-92a-1*; Extended Data Fig. 1a), known to be involved in a wide range of developmental and pathogenic processes^17^. In the adult testis, *miR-17∼92* maintains transcriptomic levels within normal values^18^ ^19^ and, in cooperation with its paralog cluster *miR-106b∼25,* it is required to sustain male fertility^20^. Interestingly, *miR-17∼92* cluster members are expressed in mouse gonads from both sexes during and after the sex determination stage (Extended Data Fig. 1b) and their predicted targets include numerous sex-related genes (see below). However, a possible role of the miR*-17∼92* cluster in sex determination has not been previously investigated.

**Figure 1.**
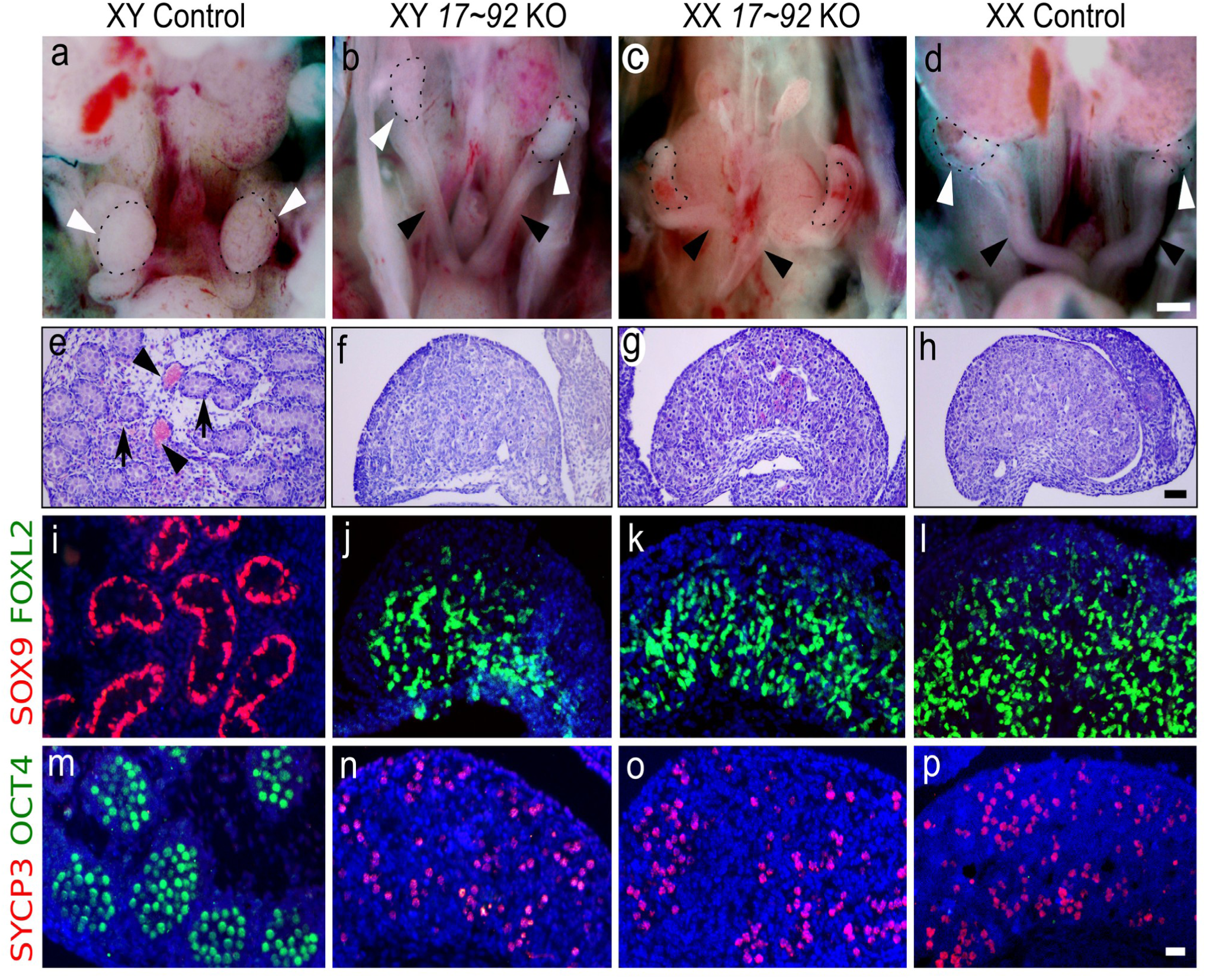
Male-to-female sex reversal in *miR-17∼92* KO mice. **a-d**, Urogenital anatomy of control and mutant XY and XX embryos at E17.5. Gonads are encircled by a dotted line and marked by white arrowheads. Uterine cornua are marked with black arrowheads. Note that only control XY embryos develop testes, while the rest develop ovaries and uterine cornua. **e-h**, Gonadal histology at E17.5. Control testes contained solid cords (arrows), large blood vessels (arrowheads) (e). In contrast, XY mutant gonads (f) showed a typical ovarian histology lacking both cords and vessels, as observed in XX gonads (g, h). **i-l**, Double immunofluorescence at E17.5 showing that only XY control testis expressed the testicular marker SOX9 (i), while the other three genotypes expressed the ovarian marker FOXL2 (j-l). **m-p**, Double immunofluorescence for the pluripotency marker OCT4 and the meiotic prophase cell marker SYCP3 at E17.5. Only germ cells in XY control testes expressed OCT4 and showed no SYCP3 expression, evidencing the absence of meiotic cells in embryonic testes (m). Contrarily, all germ cells of the other three genotypes had entered meiosis, as they expressed SYCP3, evidencing full ovarian development (n-p). Scale bars shown in d, h, and p represent 500 μm for a-d, 100 μm for e-h; and 20 μm for i-p.

Here we demonstrate that the ablation of the *miR-17∼92* cluster, prior to sex determination, causes complete male-to-female sex reversal in mice. By combining bulk and single-cell RNA-seq (scRNA-seq) and time-course expression analyses, we demonstrate that *Sry* expression is delayed in the absence of *miR-17∼92*. As a consequence of this heterochrony, pre-supporting cells undergo a transient state of sexual ambiguity that results in failed differentiation into Sertoli cells and the subsequent activation of the ovarian program. Furthermore, we show that *miR-17∼92* ablation results in a misregulation of hundreds of target genes, simultaneously affecting the regulation of critical gene networks. Our results uncover a novel and unexpected role for miRNAs in controlling the timing of *Sry* expression and Sertoli cell differentiation.

## RESULTS

### XY embryos lacking *miR-17∼92* develop as phenotypic females

To investigate the potential role of the *miR-17∼92* cluster in mammalian sex determination, we analysed the genito-urinary system of knockout (KO) mice, generated by using the Cre/LoxP system. As *miR-17∼92* KO mice undergo neonatal lethality^21^, we focused our analysis on the embryonic day 17.5 (E17.5), when sex is already differentiated. While XY control foetuses displayed normal descended testes, XY mutants developed as phenotypic females with two uterine horns and ovaries directly inferolateral to the kidneys, similar to those observed in both XX mutants and control females (Fig. 1a-d). Histological analyses on XY KO gonads revealed a complete absence of male-specific structures, such as testicular cords. Instead, gonadal structure was completely indistinguishable from XX mutants or control ovaries (Fig. 1e-h). Double immunofluorescence showed that the testicular marker SOX9 was exclusively present in XY control gonads, whereas XY mutant gonads, as XX controls, only expressed the ovarian marker FOXL2 (Fig. 1i-l). Consistently, the activation of germ cell meiosis, an early sign of ovarian development, was observed in XY mutant and XX gonads, but not in XY controls (Fig. 1m-p). Altogether, these results show that, in the absence of the *miR-17∼92* cluster, XY mice undergo complete primary sex reversal and develop as phenotypic females.

### *miR-17∼92* regulates gonadal growth before sex determination

To explore the molecular mechanisms associated to sex reversal in *miR-17∼92* mutants, we performed bulk RNA-seq on gonads during the sex determination period and shortly afterwards (E11.5 and E12.5). A variable number of differentially expressing genes (DEGs) were identified in pairwise comparisons between the four analysed genotypes at these two stages (Extended Data Tables 1-9). The DEGs between XX and XY control gonads at E12.5 (3569 DEGs; FDR<0.05) were used as a reference set for genes with normal dimorphic expression. Log_2_-fold-change heat maps at E12.5 revealed that both XY and XX mutants acquire a female-like expression profile, with most genes displaying an ovarian-specific expression pattern (Fig. 2a). Consistent with this observation, multidimensional scale plot revealed moderate differences between mutants and controls at E11.5, with mutant conditions clustering closer together. At E12.5 these differences were more prominent, with XY control replicates clustering separately from all other samples (Extended Data Fig. 2a), indicating that mutant gonads follow the ovarian fate shortly after the sex determination period.

**Figure 2.**
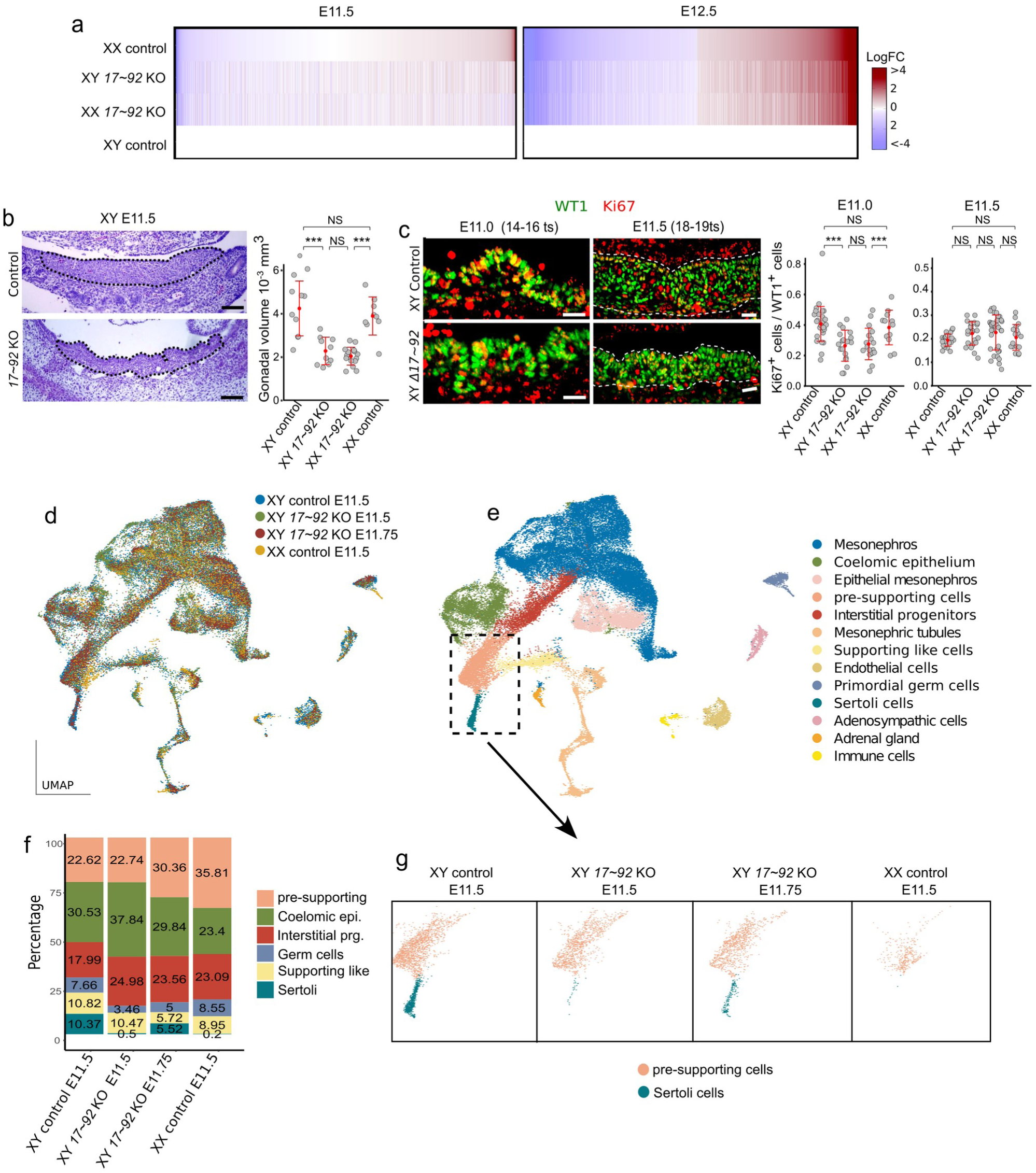
Gene expression and morphometric analysis of *miR-17∼92* KO gonads during sex determination. **a**, Log_2_-fold-change heat maps of DEGs detected between gonads of four different genotypes. Note the similarities at E11.5, as well as the female-like RNA profile of E12.5 mutants, irrespectively of their sex. **b,** Haematoxylin and Eosin staining of gonadal sections. Mutant gonads show an irregular contour, including deep transversal furrows, and are thinner and shorter than controls, with a volume reduction of 50% at E11.5 (18-19 ts). **c**, Cell proliferation analysis using double immunofluorescence for the gonadal progenitor cell marker WT1 and the cell proliferation marker, Ki67, in gonads both before (E11.0; 14-16 ts) and at the sex determination stage (E11.5; 18-19 ts). Quantification of the mitotic index (number of Ki67^+^ cells/number of WT1^+^ cells) indicate reduced cell proliferation in XX and XY mutant gonads compared to controls before (E11.0; 14-16 ts) but not at the sex determination stage (E11.5; 18-19 ts). **d-g,** single-cell RNA (scRNA-seq) analysis. For clustering and identification of populations, cells were coloured by genotypes (**d**) and by cell clusters (**e**). All major gonadal cell populations were present in both mutant and control gonads, except Sertoli cells that appeared only in XY controls at E11.5. Note that these cells were identified in XY mutants at a later timepoint E11.75 ( **f** and **g**). *, p <0.05, **, p<0.01, ***, p<0.001. Scale bars in b and c represent 50 and 20 μm, respectively.

We observed that the number of DEGs between XX and XY mutant gonads was marginal in both E11.5 and E12.5 stages (12 and 21 DEGs, respectively; Extended Data Tables 1-9), denoting the existence of *miR-17∼92*-related signatures that are shared between sexes. Furthermore, Gene Ontology analysis identified several common GO terms between E11.5 mutant and controls, in both XX and XY backgrounds (Extended Data Fig. 2b; Extended Data Tables 10 and 11). These observations suggest that *miR-17∼92* may control some non-sex related aspects of gonadal development. In particular, we found GO terms related to cell proliferation and molecular pathways involved in its control. Consistently, *miR-17∼92* KO gonads were smaller than their wildtype counterparts at E11.5, displaying a mean volume of ∼50% compared to controls (Fig. 2b). Based on these results, we investigated proliferation rates in developing gonads using double immunofluorescence against the Wilms tumor protein (WT1), a progenitor cell marker, as well as the proliferation marker Ki67. This analysis revealed that the genital ridge of mutant gonads contained less proliferating progenitor cells (WT1^+^, Ki67^+^) than that of controls, before (E11.0; 14-16 tail somites, ts) but not at the sex determination stage (E11.5; 18-19 ts; Fig. 2c). In addition, we found no differences in apoptosis in any of the stages analysed (Extended Data Fig. 2c), with the number of apoptotic cells being very low in all cases. Altogether, these results indicate that *miR-17∼92* regulates molecular pathways necessary for the normal growth of the bipotential genital ridges before the sex determination stage.

### *miR-17∼92* is required for proper *Sry* expression timing and Sertoli cell differentiation

The reduced proliferation in early *miR-17∼92* KO gonads, which is consistent with their overall reduced size, may alter the relative proportions of gonadal cell types. Therefore, we explored changes in cellular composition and identity in *miR-17∼92* KO gonads, by performing single-cell RNA-seq (scRNA-seq) experiments. We employed CRISPR/Cas9 to generate a homozygous deletion of the miRNA cluster in mouse embryonic stem cells (mESC) and subsequently derived mutant animals via tetraploid complementation assays^22^ ^23^. Of note, these mutants displayed an identical phenotype to those previously generated via Cre-Lox breeding. The profiled cells were clustered based on their transcriptional similarities, recapitulating all major cell populations previously described in developing gonads^24^ (Fig. 2d and e and Extended Data Fig. 3a and b). All cell clusters were present in mutant and control samples, irrespectively of the sex, with the only exception of Sertoli cells, a male-specific cell type that only appeared in XY control gonads (Fig. 2f and g). To elucidate if the lack of Sertoli cells in XY *miR-17∼92* mutant gonads was due to delayed or impaired testis differentiation, we also analysed an additional sample of XY *miR-17∼92* KO gonads at a later stage (E11.75). Cluster integration confirmed that Sertoli cells were also present in *miR-17∼92* mutants, albeit their number was lower than in XY control gonads at E11.5 (5.52% and 10.37%, respectively; Fig. 2f and g). These results indicate that *miR-17∼92* is required for proper extent and timing of Sertoli cell differentiation.

**Figure 3.**
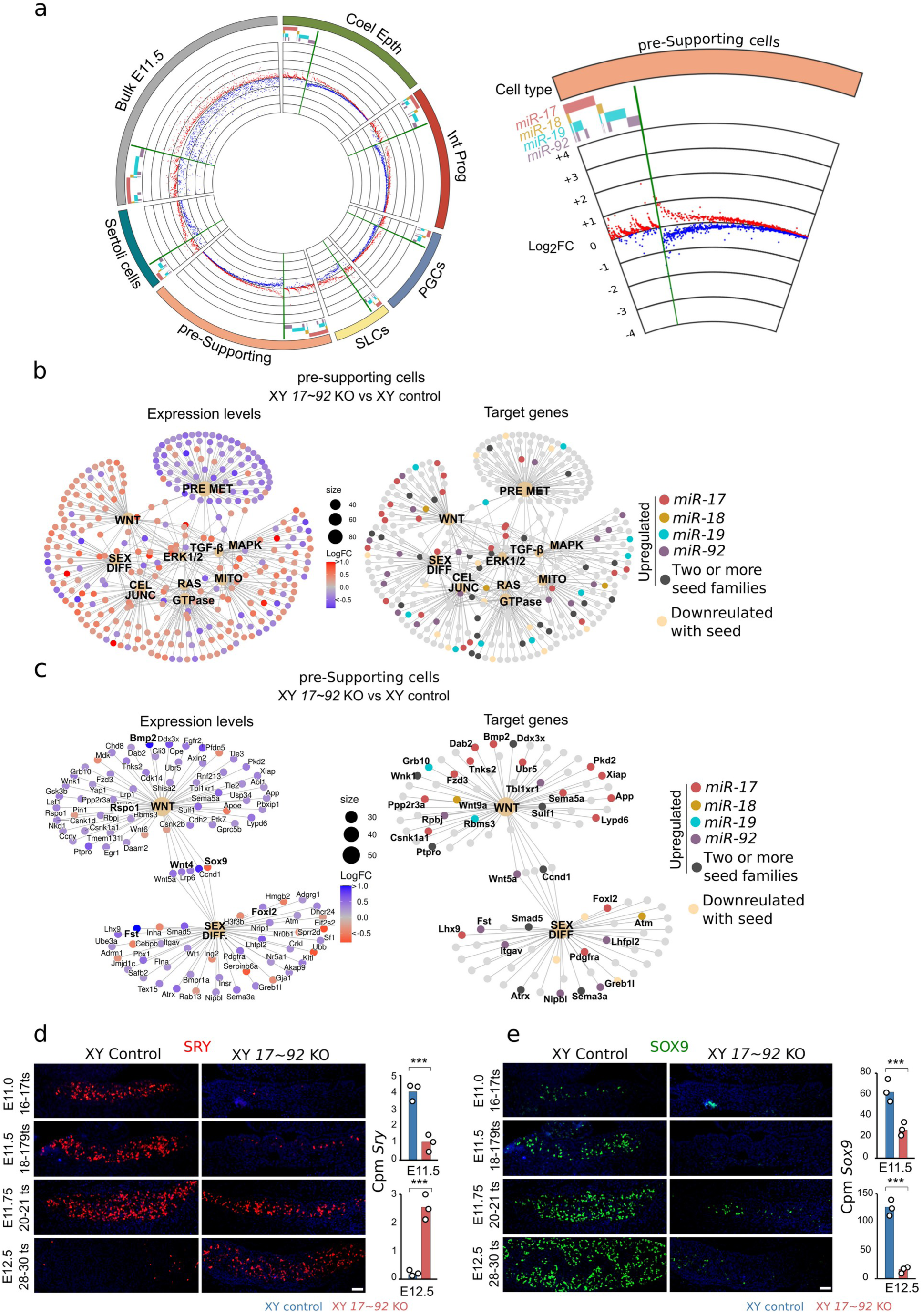
Effects of *miR-17∼92* cluster depletion on gene expression. **a,** Circos plot of DEGs in cell populations of XY *miR-17∼92* KO gonads (see Extended Data Fig. 4 for a high resolution image). Left panel, genes with predicted binding sites for one or more *miR-17∼92* seed families (target genes) are grouped at the left side of each sector and marked with a coloured bar in the outer circle (separated by a green bar). A scatter plot of log2FC is shown in the inner circle, with up-and downregulated genes labelled in red and blue, respectively. Right panel, magnification of the pre-supporting cells sector. Non-target genes (on the right of the green bar) do not show any particular bias towards up-or downregulation, whereas *miR-17∼92* target genes (on the left) are preferentially upregulated. **b,** Gene-concept networks using DEGs between XY mutant and control pre-supporting cells at E11.5-E11.75. Log_2F_Cs (at left) and predicted targets for the four *miR-17∼92* seed families (at right) are depicted. The main deregulated functions and cell signalling pathways are shown. **c,** Gene-concept network focusing on DEGs which special relevance in sex determination. Log_2_FCs (at left) and predicted targets for the four *miR-17∼92* seed families (at right) are depicted. Most genes of the WNT signalling pathway are downregulated, including several genes carrying *miR-17∼92* target sequences. Note the deregulation of important genes for sex determination, such as *Sox9*, *Foxl2* and *Wnt4*. **d,** Immunofluorescence for SRY on control and mutant gonads during sex determination. Note that SRY expression is delayed in XY mutant gonads, compared to XY controls. *Sry* expression levels, calculated using the CPMs of the bulk RNA-seq analysis, are shown on the right panel. **e**, Immunofluorescence for SOX9 on control and mutant gonads during sex determination. Note that the onset of SOX9 expression is delayed in mutant gonads (starts at 20-21 ts), and the number of SOX9^+^ cells is reduced. *Sox9* expression levels are also shown on the right panel. Coel. Epth., Coelomic Epithelium; Int. Prog., Interstitial Progenitors; PGCs, Primordial Germ Cells; SLCs, supporting-like cells. Scale bars shown in the last picture of d and e represent 50 μm for all the pictures. ***, p<0.001.

To investigate the mechanisms of *miR-17∼92-*mediated gene expression, we examined the DEGs between XY control and mutant gonads (Extended Data Tables 12-18). Although miRNAs mostly control protein production, previous studies have shown that the changes in protein levels induced by miRNAs are largely mirrored at the transcript level^25^ ^26^ ^27^. Since the individual components of the *miR-17∼92* cluster can be grouped into four “seed families”, based on their main target sequences^17^ (Extended Data Fig. 1a), we generated cumulative distribution fraction (CDF) plots of the DEG Log_2_FCs. In both bulk and sc-RNAseq datasets, the predicted target genes of the four seed families were preferentially upregulated in mutant gonads, compared to non-target genes (Extended Data Fig. 3c), indicating that they are negatively regulated by *miR-17∼92* during normal testis development. To visualize the relative contribution of each seed family to gene regulation, we generated circos plots in which the direction and amplitude of deregulation, as well as the presence of predicted binding sites, was depicted (Fig. 3a; Extended Data Fig. 4). As previously noted, the predicted target genes of the four seed families were preferentially upregulated in the XY mutant gonads compared to XY controls. However, we observed that many non-target genes of the cluster were also deregulated, although in this case, there was no preference in the direction of deregulation. Furthermore, the changes in the expression levels of the deregulated genes were generally modest (less than two times; Fig. 3a and Extended Data Tables 12-18). These results are consistent with previous studies in which the cluster was conditionally deleted in mouse embryonic tail bud and heart, which showed that *miR-17∼92* members act as fine-tuners of large gene networks rather than regulating the expression of a few genes coding for specific transcription factors^28^.

**Figure 4.**
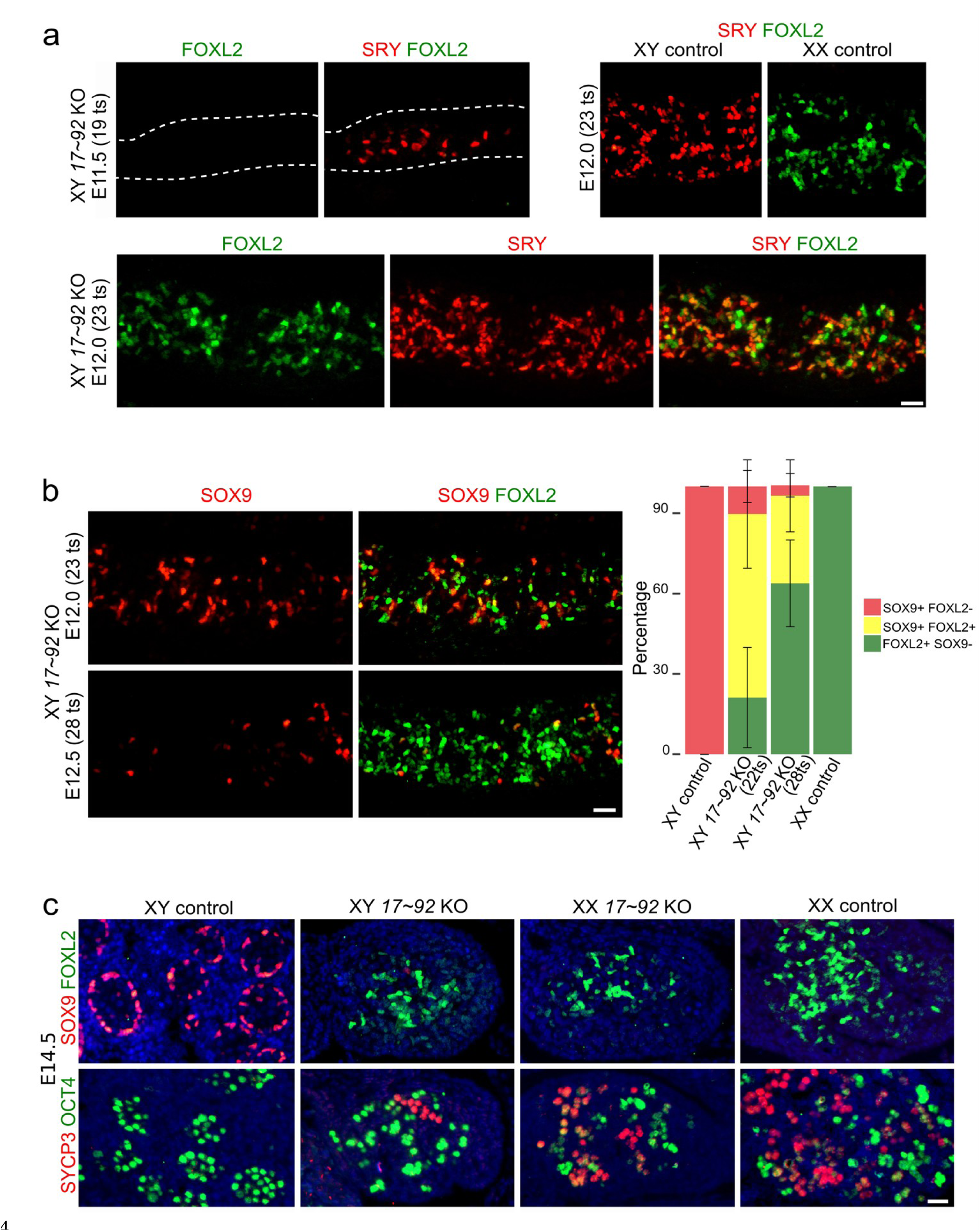
Time course of granulosa and oocyte cell differentiation in XY *miR-17∼92* KO gonads. **a,** Double immunofluorescence for SRY and FOXL2 in gonads during (E11.5) and shortly after the sex determination stage (E12.0). **b**, Double immunofluorescence for SOX9 and FOXL2 after sex determination. Cells showing colocalization of both proteins are common at E12.0 but very scarce at E12.5. **c**, Double immunofluorescences for SOX9 and FOXL2 (upper panel) and for SYCP3 and OCT4 (lower panel) in gonads at E14.5. SOX9^+^/FOXL2^+^ cells are almost completely absent at this stage, in which only FOXL2 expression persists. Meiotic oocytes (SYCP3^+^ cells) are seen in XY mutant gonads, evidencing complete ovarian differentiation. The scale bars shown in the lower right images of panels a, b and c represent 25 μm for a and b, and 20 μm for c, respectively.

Since Sertoli cells do not differentiate properly in XY mutant gonads, we explored the effects of *miR-17∼92* ablation on their progenitors, the pre-supporting cells. Gene Ontology analysis on DEGs from XY control and mutant pre-supporting cells revealed an enrichment in categories associated with known biological proccess, (“*cell junction assembly*”^29^, “precursor metabolite energy”^30^, and “reproductive system development”) and signalling pathways (MAPK^31^, WNT^32^ ^7^) operating in gonadal supporting cell differentiation. (Extended Data Fig. 5a; Extended Data Table 19). Gene concept analysis also revealed a large network in which different sub-networks shared numerous genes and *miR-17∼92* targets (Fig. 3b). In general, the percentage of upregulated *miR-17∼92* target genes in XY mutant pre-supporting cells for each GO category ranged from 30% to 40%, whereas downregulated target genes where normally below 7% (Extended Data Fig. 5b). Again, these results suggest that the *miR-17∼92* cluster regulates the expression of numerous target genes that may, in turn, fine-tune the expression of large gene networks during sex determination. Among these gene networks we found “*sex differentiation*” and “*Wnt signaling pathway*”, the latter being essential for ovary differentiation^32^ ^7^. In particular, DEGs belonging to the WNT pathway were preferentially upregulated in XY mutant pre-supporting cells (52 genes upregulated and 7 downregulated; Fig. 3c), and 40% of these genes were putative targets of the *miR-17∼92* cluster (Fig. 3c; Extended Data Fig. 5b). In addition, *Foxl2* and *Fst*, two key factors during ovarian development^33^ ^34^, which are also *miR-17∼92* targets, were upregulated in XY mutant gonads (Fig. 3c; Extended Data Table 14). These results suggest a function for the *miR-17∼92* cluster in maintaining female-promoting genes and signalling pathways downregulated during early testis differentiation.

However, we also found that *Sox9,* which encodes a transcription factor essential for testis development but is not a putative *miR-17∼92* target, was downregulated in mutant pre-supporting cells (Fig. 3c; Extended Data Table 14). Therefore, we investigated the expression of the two key factors necessary for Sertoli cell differentiation in mammals: SOX9 itself, and its main activator, the testis-determining factor SRY. Immunofluorescence analyses revealed that cells expressing the SRY protein were already present by E11.25 in XY control testes (16-17 ts), with their number peaking at E11.5-E11.75 (∼18-21 ts) and declining afterwards until almost disappearing by E12.5 (28-30 ts). In contrast, the first SRY^+^ cells in XY mutant gonads were detected at E11.5 (18-19 ts), and this number progressively increased until E12.5 (28-30 ts). Consistently, *Sry* transcript levels in mutant XY gonads were significantly lower than those of XY controls at E11.5, whereas the opposite situation was observed at E12.5. (Fig. 3d; Extended Data Tables 3 and 7). As previously described^35^, the number of SOX9^+^ cells increased dramatically in E11.5 control testes and remained elevated at subsequent stages. In XY mutants, however, SOX9 was not detected at E11.5 (18-19 ts), and SOX9^+^ cells were notably less abundant than in XY controls at subsequently stages (20-21ts and onwards), as shown also at the transcript level (Fig. 3e). It is known that *Sry* must act within a critical time window during mouse sex determination stage to induce testis differentiation^36^ ^37^. Thus, the temporal delay of *Sry* expression and the reduced number of SRY-expressing cells provides a mechanistic basis for the deficient *Sox9* upregulation and Sertoli cell differentiation of XY *miR-17∼92* KO gonads.

### A transient state of sexual ambiguity in *miR-17∼92* KO pre-supporting cells

We next investigated granulosa cell differentiation by immunofluorescence for the ovarian marker FOXL2. In XY mutant gonads, we observed that FOXL2 is activated shortly after the sex determination window (E11.75; ∼20-21 ts), displaying a sustained expression throughout all subsequent stages (Extended Data Fig. 6). This expression onset coincides with that observed in XX control gonads and occurs after the initiation of *Sry* expression (Fig. 4a). This profile is consistent with the hypothesis that granulosa cell differentiation in XY mutant gonads results from delayed *Sry* and *Sox9* expression, although a precocious activation of female-promoting genes carrying *miR-17∼92* binding sites cannot be completely ruled out. The analysis of gonads at later stages showed that the vast majority of XY mutant cells expressing FOXL2 did also express SRY (Fig. 4a), revealing that *Foxl2* was preferentially upregulated in pre-supporting cells. Consistently, FOXL2-SOX9 double immunofluorescence showed that, at E12.0 (22-23ts), most of FOXL2^+^ cells coexpress also SOX9^+^ (Fig. 4b). Nevertheless, XY *miR-17∼92* KO gonads also contained many FOXL2^+^ SOX9^-^ cells, both at E12.0 and E12.5, suggesting that many granulosa cells in XY *miR-17∼92* KO gonads differentiate directly from undifferentiated pre-supporting cells. These results indicate that mutant pre-supporting cells undergo a transient state of sex ambiguity, characterized by the concomitant initiation of both the female and the male pathways.

Previous studies have shown that overexpression of *Foxl2* during early testis development can cause male-to-female sex reversal^38^ ^39^. Thus, differentiation of FOXL2^+^ granulosa cells at early stages of gonadal development appears to be a key event underlying *miR-17∼92-*associated sex reversal. Interestingly, *Foxl2* carries a functionally validated *miR-17* binding site in its 3’UTR^40^ (Extended Data Fig. 7a) suggesting that early *Foxl2* upregulation in XY mutant gonads could be a direct consequence of *miR-17∼92* ablation. To test this hypothesis, we deleted the *miR-17* binding site from the 3’ UTR of *Foxl2* in XY mouse embryonic stem cells (mESC), and generated E11.5 and E12.5 embryos via tetraploid complementation assay (XY *Foxl2-3’-UTR^del/del^*). The absence of the *miR-17* binding site did not affect *Foxl2* transcript stability in eye (Extended Data Fig. 7b). However, XY mutant embryos exhibited normal testis differentiation, with no evidence of *Foxl2* upregulation neither at the transcript nor at the protein level (Extended Data Fig. 7c; Extended Data Tables 20 and 21). Therefore, the upregulation of *Foxl2* at E12.0 in XY *miR-17∼92* mutants is mostly an indirect effect derived from events occurring at earlier stages, likely associated to the low *Sox9* expression levels that are insufficient to antagonize the female pathway. In fact, *miR-17∼92* ablation in Sertoli cells after the sex determination stage does not result in *Foxl2* upregulation^18^. Nevertheless, a transient down-regulation of *Foxl2* by *miR-17∼92* in the testis shortly after the sex determination period cannot be completely excluded.

In addition to *Foxl2*, other female-specific markers, including *Wnt4*, *Rspo1*, *Bmp2* and *Fst* were upregulated in XY mutant gonads at E12.5, whereas male-specific markers such as *Ptgds*, *Sox8*, *Amh,* and *Dhh* were downregulated (Extended data Fig. 8a; Extended Data Tables 3 and 7). This indicates that the dual sexual program observed in XY mutant gonads shortly after sex determination is resolved at later stages in favour of the female pathway. At E14.5, only a few scattered SOX9^+^ cells remain in XY mutant gonads, which showed a normal ovarian development including the presence of FOXL2^+^ supporting cells, meiotic germ cells and a lack of male-specific cell populations or structures, evidencing complete male-to-female primary sex reversal (Fig. 4c; Extended Data Fig. 8b).

## DISCUSSION

Since miRNAs discovery, numerous attempts have been made to elucidate their role during sex determination. However, to date, none of them could demonstrate that such post-transcriptional regulators can play a fundamental role in this process^9^ ^6^. Indeed, previous studies compromising the biogenesis of these molecules suggested that they are dispensable for sex determination^15^ ^16^. Contrarily, here we show that a single cluster of miRNAs is essential for mouse testis differentiation. Our results reveal that *miR-17∼92* controls two main events of mouse sex determination, namely early gonadal growth and *Sry* expression dynamics. Proliferative growth of the bipotential gonadal primordia is essential to establish the pool of gonadal somatic progenitors, including pre-Sertoli cells, whose number must reach a minimum threshold prior to sex determination to ensure subsequent testis differentiation^41^ ^42^. The gonads of both XX and XY *miR-17∼92* KO embryos are smaller than those of control mice (50% reduction) at the sex determination stage, which is a consequence of reduced proliferation during the bipotential stage. This proliferative reduction in mutant gonads appears to affect all progenitor cell types similarly, as their relative numbers do not differ significantly from those of controls, notably in the case of the pre-supporting cell lineage (Fig. 2f). Hence, the number of pre-Sertoli cells present at the time of sex determination (E11.5), although reduced, should be sufficient to initiate testis differentiation^42^. Despite of this, *miR-17∼92* mutant gonads contain almost no Sertoli cells at E11.5, and shortly later (E11.75) their number is only half that of control testes, a fact that compromises testicular but not ovarian differentiation. The lack of Sertoli cells in XY *miR-17∼92* mutant gonads at the sex determination stage is associated with the heterochronic expression of *Sry*, which must act within a critical time window to activate *Sox9* and trigger the male pathway^36^ ^37^. *Sry* expression in XY *miR-17∼92* mutants is delayed by about 12 hours, which explains the lack of *Sox9* up-regulation during sex determination and the subsequent XY sex reversal. Interestingly, miRNA knockouts have been repeatedly reported to cause heterochronic shifts in developmental processes^10^ ^11^ ^12^ across a wide range of animal models. In fact, the miRNAs of the *miR-17∼92* cluster are evolutionarily well conserved, appearing around 500 Mya in primitive vertebrates, where they have been shown to regulate genes involved in gonadal development and sex change^43^. Overall, this suggests the existence of mechanisms for mRNA function that might be commonly shared across metazoan evolution.

The delayed *Sry* expression is likely indirectly derived from the lack of *miR-17∼92,* as *Sry* itself is not a predicted target of these miRNAs. Our results also show that *miR-17∼92* targets were preferentially upregulated in XY mutant pre-supporting cells during sex determination, with a misregulation of large gene networks involved in *Sry* regulation and Sertoli cell differentiation. One of these networks is the MAPK signalling pathway, which is involved in the control of both gonadal cell proliferation and *Sry* expression timing. In fact, similar cases of XY sex reversal were described in mice with disrupted MAPK signalling^31^ ^44^ ^45^. We found that 27 genes of the MAPK pathway were deregulated in XY *miR-17∼92* KO pre-supporting cells, 40% of them being upregulated target genes (Extended Data Fig. 5a and b and 9a). Interestingly, *Gadd45γ,* which is required for pre-Sertoli cell fate specification *in vivo* by promoting p38 MAPK signalling^45^, shows reduced expression in mutant gonads (Extended Data Fig. 9a and b). In addition, *miR-17∼92* regulates gene networks involved in ovarian differentiation in gonadal pre-supporting cells (Fig. 3b). One of these is the WNT signalling pathway, which antagonizes Sertoli cell differentiation and is upregulated in *miR-17∼92* XY mutants (Fig. 3c), as expected in gonads that eventually develop as ovaries. Upregulation of the WNT signalling pathway is probably an indirect effect of the inactivation of the male genetic program, but its modest amplitude at E11.5 and the high number of *miR-17∼92* putative targets genes that are upregulated in this pathway (40 %) also argue for a complementary direct effect of *miR-17∼92* depletion, at least during the initial stages of sex determination. Hence, an additional function of *miR-17∼92* could be to modestly negatively regulate the WNT signalling pathway at the time of sex determination, an effect that favours testis differentiation but does not affect ovarian development. Another biological process affected in XY mutant cells is “*generation of precursor metabolites and energy*”, which is downregulated in XY mutant pre-supporting cells (Fig. 3b and Extended Data Fig. 5). Interestingly, it was previously shown that a high-glucose metabolic state is required in developing pre-supporting cells for the establishment of SOX9 expression and testis differentiation^30^.

Taken together, our results show that *miR-17∼92* acts as a cross-cutting regulator that modulates the expression of genes networks and signalling pathways that control processes including proliferative growth, energy metabolism, and female pathway antagonism. As such, *miR-17∼92* primes the bipotential gonadal primordium to ensure proper *Sry* expression timing and subsequent testicular differentiation. In summary, our study uncovers a novel role for the *miR-17∼92* cluster and for miRNAs in controlling the early steps of mammalian sex determination.

## METHODS

### Transgenic mice

To generate mice with a null deletion of the *miR-17∼92* cluster we crossed mice with a floxed allele of *miR-17-92* on a C57BL/6 × 129S4/SvJae background acquired at the Jackson Laboratory (Bar Harbor, ME, USA; Stock 008458)^17^ with *Tg(CAG-cre)^1Nagy^* mice (MGI:3586452^46^). The resulting *Tg(CAG-cre)^1Nagy^;miR-17∼92^+/del^* offspring was backcrossed to *miR-17∼92^flox/flox^* mice to obtain *Tg(CAG-cre)^1Nagy^;miR-17∼92^del/del^* embryos. Extended Data Fig. 10 shows the effective deletion of the *miR-17∼92* region in mutant mice. For timed pregnancies, plugs were checked every morning after mating and noon was taken as embryonic day (E) 0.5. Genotyping was carried out on genomic DNA derived from adult tails or embryonic yolk sacs using previously described primers^21^ ^47^. To generate mice with a deletion of the *miR-17* seed sequence in the 3’ UTR of *Foxl2* (*Foxl2-3’-UTR^del/del^*) we used the CRISPR/Cas9 technology combined with tetraploid complementation assays for embryo generation^22^ ^23^. The single guide RNA (sgRNA) was designed with Benchling (https://www.benchling.com/crispr/) and cloned into the CRISPR/Cas9 pX459v2 plasmid (ref. 62988 Addgene). Mouse embryonic stem cells (C57BL/6J-129 F1 hybrid background, G4 mESC) were cultured on a layer of CD1 fibroblast and transfected with the sgRNA using FuGENE HD (Promega, Madison, WI, USA). Clones were picked, genotyped and expanded as described previously^22^. Clones homozygous for the deletion were selected for tetraploid aggregation. Primers used for genotyping were as follows: 5’-GAAATCCCGTGACCT-GGTGG–3’ and 5’-GCTAGCTGCTGAACTACGGT-3’. The sgRNA sequence was 5’-CACCGTTGTGTTTGTACGTGTGTG-3’.

### Histological and immunostaining methods

For histology, embryos were dissected, collected in PBS, fixed in Serrás solution (ethanol : 37% formaldehyde : acetic acid, in proportions 6 : 3 : 1, respectively), embedded in paraffin, sectioned (5μm thick), and stained with haematoxylin and eosin. For double immunofluorescence, the two primary antibodies (raised in two different mammalian species) were incubated overnight at 4°C and then appropriated conjugated secondary antibodies were applied. Used antibodies are listed in Extended Data Table 22. Digital photomicrographs were taken in a Nikon Eclipse Ti microscope (Nikon Corporation, Tokio, Japan).

### Morphometric analyses

The shape of the embryonic gonad is near cylindrical but has an irregular surface, including deep furrows, mainly in mutant embryos. Hence, its volume could not be accurately calculated using the formula of a cylinder. Instead, we measured the area of the gonadal image in a 10× photomicrograph taken from a medial (that showing the maximum gonadal area), longitudinal section. For this, we used the *ImageJ*application (https://imagej.nih.gov/ij/) set at a scale of 2940 pixels per mm (the value corresponding to images taken using a 10× objective). The contour of the gonad was encircled with a line using the *freehand selection* tool and its area (A) was measured in square mm (this operation was repeated twice for each gonad and the mean value was used for further calculations). Similarly, the maximum length (L) of the gonad was depicted with a line and measured in mm. The gonadal volume (V) was calculated using the formula: V = (πA²)/(4L). The volume of the two gonads was calculated in 4 control males, 2 control females, 3 mutant males, and 4 mutant females at the 18-19 ts stage.

The mitotic index of the coelomic epithelium-derived cells was calculated by dividing the number of proliferating cells (Ki67^+^) by the total number of coelomic epithelium cells (WT1^+^) present in gonadal images photographed using a 20× objective. For this, double immunofluorescence preparations were performed on longitudinal sections of embryonic gonads from embryos both at the 14-16 ts stage (3 control males, 3 control females, 4 mutant males, and 3 mutant females) and at the 18-19 ts stage (4 control males, 2 control females, 3 mutant males, and 4 mutant females).

### Analysis of apoptosis

Apoptotic cells present in gonadal histological preparations were revealed by the terminal deoxyribonucleotidyl transferase (TDT)-mediated dUTP-digoxigenin nick end labelling (TUNEL) assay using the *Fluorescent* In Situ *Cell Death Detection Kit* (Roche ref. 11684795910, Roche, Mannheim, Germany), according to the manufacturer’s instructions. We omitted the enzyme solution for negative controls.

### Bulk RNA-seq

For RNA-seq, total RNA was purified from gonad-mesonephos complexes at E11.5 and from isolated gonads at E12.5 using the RNeasy Micro Kit (Qiagen, Hilden, Germany). For each genotype, 3 replicates were used, including 2 pair of gonads per replicate. Libraries were prepared with the NEBNext Ultra II Directional RNA Library Prep Kit for Illumina (New England Biolabs, Ipswich, MA, USA; E11.5 samples) and the TruSeq stranded mRNA Library prep (Illumina, San Diego, CA, USA; E12.5 samples) and sequenced with the Illumina Hiseq4000 platform (2×75 PE).

### Single cell RNA-seq

Three to four pairs of gonads per genotype, sex and stage were pooled and individual cell suspensions were performed using 0.05% Trypsin-EDTA (ref. 59417C, Sigma) for 7 minutes at 37°C with pipetting every 2-3 minutes. Trypsin digestion was stopped by adding 5% BSA (ref. B9000S, New England Biolabs) and the cell suspension was passed through a 40 µm Flowmi cell strainer (ref. 15342931, Fisher scientific). Finally, cells were collected by centrifugation (300g for 5 min at 4 °C) and the pellet was resuspended in PBS for cell counting and viability check and then fixed with methanol (98%) added dropwise. Fixed cells were kept on ice for 15 min. and subsequently moved to −80°C for extended preservation. After rehydration, about 10,000-20,000 cells were loaded on a 10x Chromium controller and sc-RNA libraries were prepared using the Chromium Next Gem Single Cell 3’ Reagent Kit v3.1 (PN-1000128) following the manufacturer’s instructions. Finally, libraries were sequenced using Illumina NovaSeq 6000 SP with the following parameters: paired-end, 28 + 8 + 91 cycles. The sequencing depth was 200 million reads per sample. Two technical replicates were performed per genotype except for the XY E11.75 KO mice, with one replicate.

### Bioinformatic analyses

The bulk RNA-seq reads were mapped to the mm10 mouse genome and counted with the *align* and *featureCounts* function from the R subread package^48^. Only genes with 1 or more cpm (counts per million) in at least two of the samples were considered to be expressed and used for further analysis. Analysis of differential gene expression was performed with edgeR^49^ Genes were considered to be differentially expressed at a false discovery rate (FDR) <0.05. Gene Ontology analysis was performed with the enrichGO function of the clusterProfiler bioconductor package ^50^. General terms and redundant terms were not displayed. For gene-concept analysis we used the cnetplot function of the same suite. Prediction of target sites for the *miR-17∼92* seed families was performed using TargetScan (mouse version 8)^51^. To analyse the gonadal cell lineage-specific expression of the individual members of the *miR-17∼92* cluster we used the RMA normalized data from Jameson et al.^1^ and differential expression was assessed with the limma R package.

Regarding single cell RNA-seq, each single cell library was aligned to the mm10 reference genome (mm10_v3.0.0, obtained from 10x Genomics) and transcriptome (modified to include a second exon of *Sry* as stated in previously^24^ and filtered using Cell Ranger Software; version 6.1.1) using default parameters. The filtered counts matrices of all the samples with two technical replicates were merged using Seurat (version 4.0.2)^52^ before proceeding with downstream analyses. For the male samples, once they were merged, thresholds for percentage of mitochondrial genes were set in order to filter low quality cells (Male control 11.5 = 10%, Male mutant 11.5 = 10%, Male mutant 11.75 = 9%). Also, blood cells were filtered from the analyses according to their percentage of beta globin transcripts (male control 11.5 = 1.43%, male mutant 11.5 = 0.39%, male mutant 11.75 = 0.73%). Finally, only cells with nCount_RNA < 30000 (transcripts) and nFeatures_RNA < 7000 (genes expressed) were considered. After quality check we analysed 23527, 17057, 8492, and 3669 cells from gonads of XY controls at E11.5, XY *miR-17∼92* mutants at E11.75, XY *miR-17∼92* mutants at E11.5 and XX controls at E11.5, respectively. Next, data was normalized using default settings, 2000 variable features were identified using the default Seurat method (vst). Data was then scaled regressing the nCount_RNA (transcripts) and percentage of mitochondrial genes. Finally, clusters were obtained with a resolution of 1.2 with the FindClusters function from Seurat. For the female samples, the filtering was made at this step by excluding those clusters with low quality (low nFeatures_RNA or high number of beta hemoglobin genes expressed, compared to the rest of the clusters). Doublets were excluded using DoubletFinder (version 2.0.3)^53^. In order to generate a single dataset with all samples, the Seurat objects were integrated using the Canonical Correlation analysis (CCA) method^54^ with 40 dimensions and using the 2000 most variable features. Then, the integrated object was scaled as previously stated and clusters were called at resolution 1 with the FindClusters function from Seurat. A single cluster was manually removed (it contained cells only from control males at 11.5 and was scattered along all the UMAP). Markers for each cluster were identified using the FindMarkers function of Seurat with default parameters. Clusters were annotated according to their expression markers according to previous bibliography^24^. Finally, DEG lists between the clusters and samples of interest were generated using the Model-based Analysis of Single cell Transcriptomics (MAST) method^55^. Mitochondrial and ribosomal genes as well as blood markers were manually removed from the DEG lists.

## Acknowledgements

We are indebted to the Animal Experimentation Unit of the University of Granadás Centre for Scientific Instrumentation for kindly extending the maintenance of our mouse colony temporally after the end of our financial support for this work. We thank the sequencing core, transgenic unit and animal facilities of the Max Delbrück Centre for Molecular Medicine for technical assistance. We thank R. Kühn, C. Scholl, M. Altmann, G. Kussagk, S. Bomberg, S. Reissert-Oppermann and C. Westphal for their support with the transgenic work. The authors thank Dr. Dagmar Wilhelm (University of Melbourne) for kindly providing us with the SRY antibody and Dr. James F. Martin (Baylor College of Medicine) for providing us with a *pri-miR-17∼92 in situ* probe. This work was supported by grants from the Andalusian Government, Junta de Andalucía, P20_00583 to R. Jiménez and F.J. Barrionuevo; P11-CVI-7291 to M. Burgos; and by a grant from the Deutsche Forschungsgemeinschaft (International Research Training Group 2403) to I. Mota-Gómez and D.G. Lupiáñez. The authors thank the Spanish Ministry of Science and Innovation for the PhD fellowship granted to A. Hurtado.

## Authors contributions

RJ and FJB conceived the project; RJ, FJB, DGL, and MB designed the experiments; RJ, FJB, AH, IM-G, ML, FMR, and JJ performed laboratory experiments; FJB, MB, and IM-G carried out bioinformatic analyses; RJ, FJB, MB, and DGL collected the financial support; RJ and FJB wrote the first draft of the manuscript and all authors contributed to the final version.

## The authors declare no competing interests

### Ethics declaration

All animal experiments in this study were approved by the University of Granada Ethics Committee for Animal Experimentation (exp. No: 2011–341) and by the Landesamt für Gesundheit und Soziales, Berlin, Germany under license number G0111/17, and were performed in accordance with the relevant guidelines and regulations dictated by these Committees.

**Correspondence** and requests for materials should be addressed to Darío G. Lupiáñez; Rafael Jiménez; Francisco J. Barrionuevo.

## Extended Data Figures

**Extended Data Figure 1.**
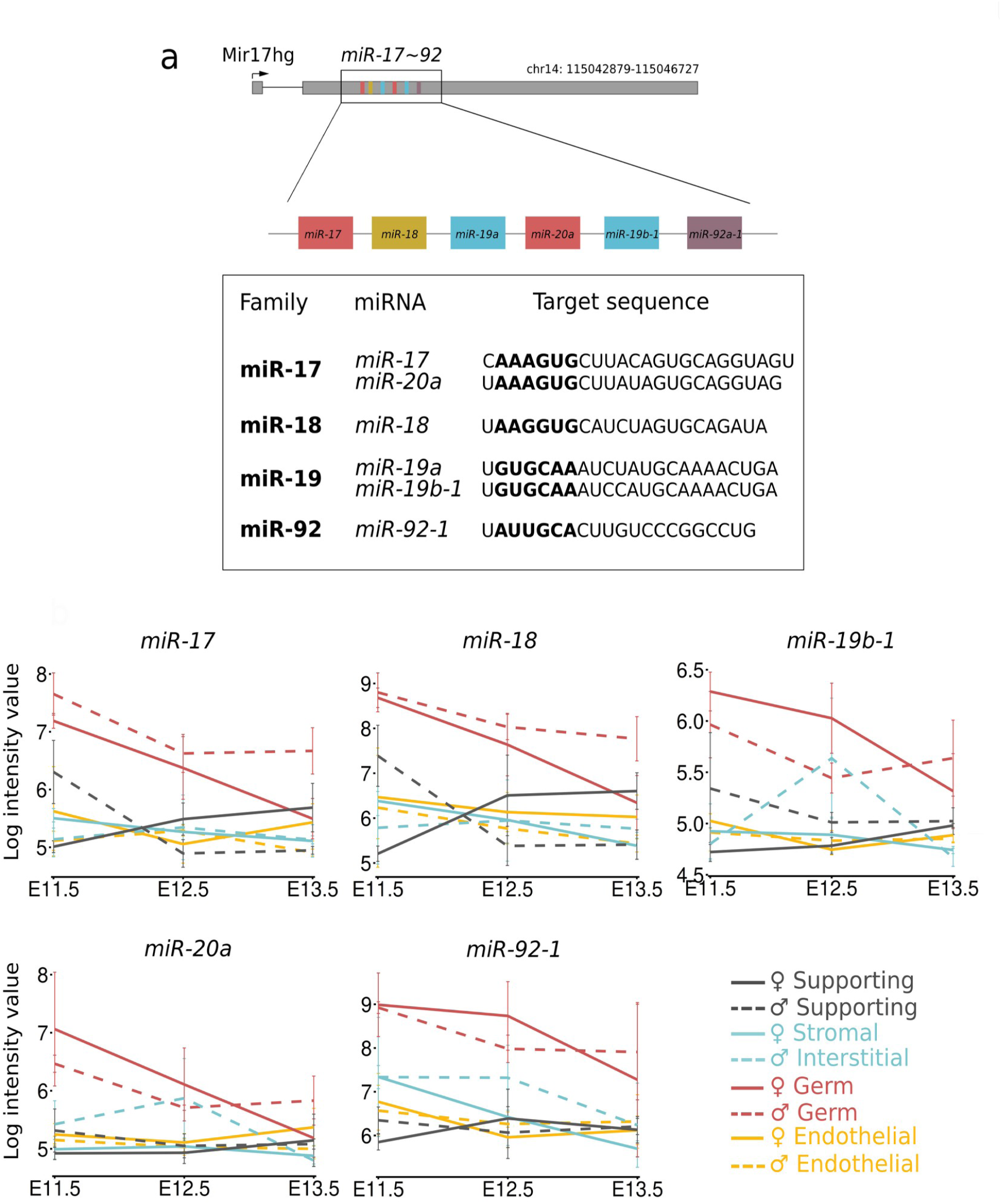
The mouse *miR-17∼92* cluster. **a**, Genomic organization of the mouse *miR-17∼92* cluster (top). The members of the *miR-17∼92* cluster are grouped in four families according to their seed sequences, the regions considered to be the most important for target selection (in bold; bottom). **b**, Expression of *miR-17∼92* members in the main gonadal cell populations during mouse sex determination (E11.5) and early gonad differentiation. Data are obtained from Jameson and colleagues^1^.

**Extended Data Figure 2.**
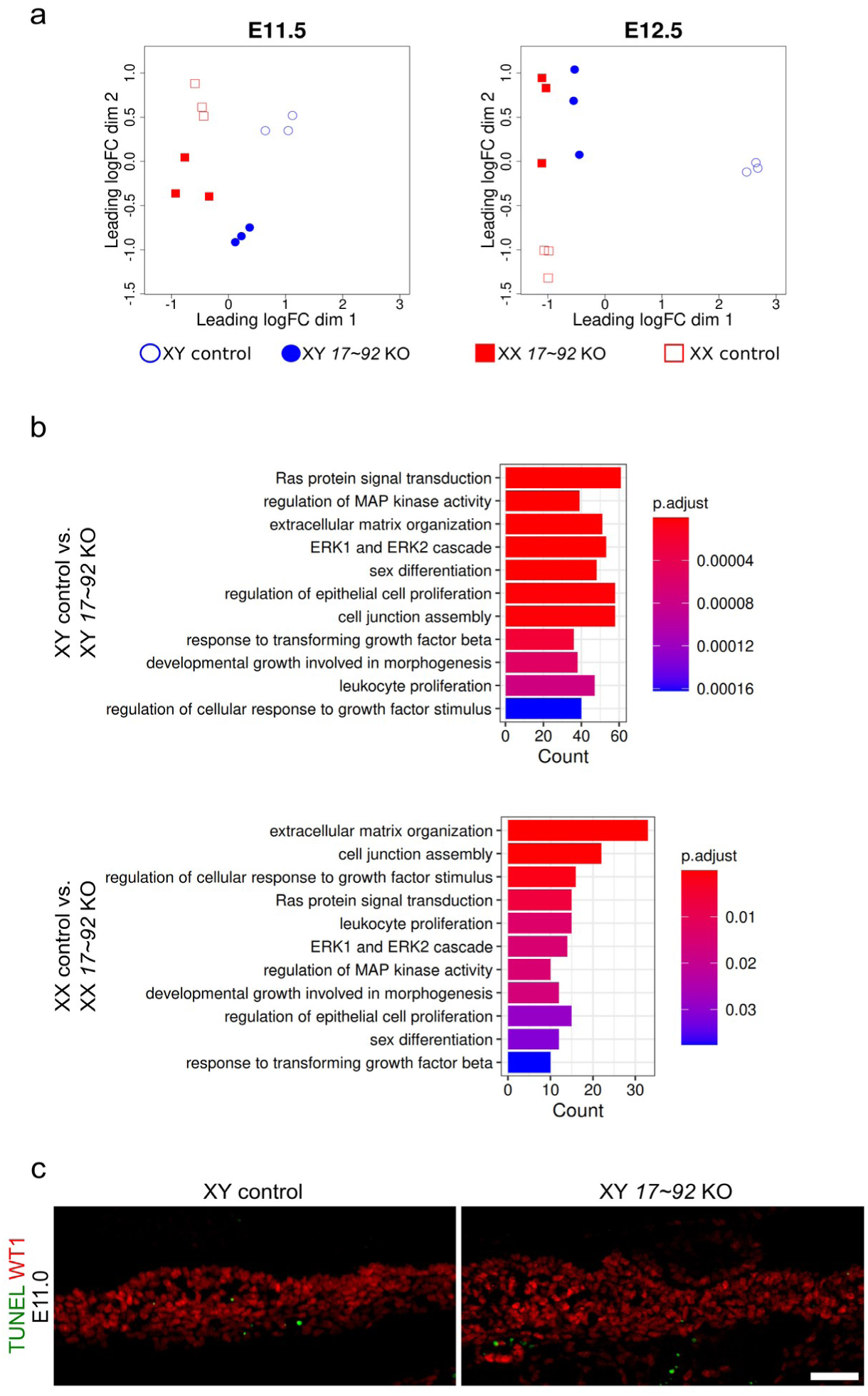
Transcriptomic analyses and apoptotic cells detection of *miR-17∼92* KO gonads. **a**, Multidimensional scaling plots of transcriptomic data from XX and XY control and *miR-17∼92* KO gonads at E11.5 and E12.5. At E11.5 replicate samples of all conditions clustered close together, whereas at E12.5 the XY control group was clearly separated from the other samples. **b**, Gene Ontology analyses revealed common deregulated biological processes between controls and mutants in both XX and XY gonads. **c,** TUNEL assay showed no difference in apoptosis between control and mutant gonads during early gonadal differentiation. Scale bar represents 50 μm.

**Extended Data Figure 3.**
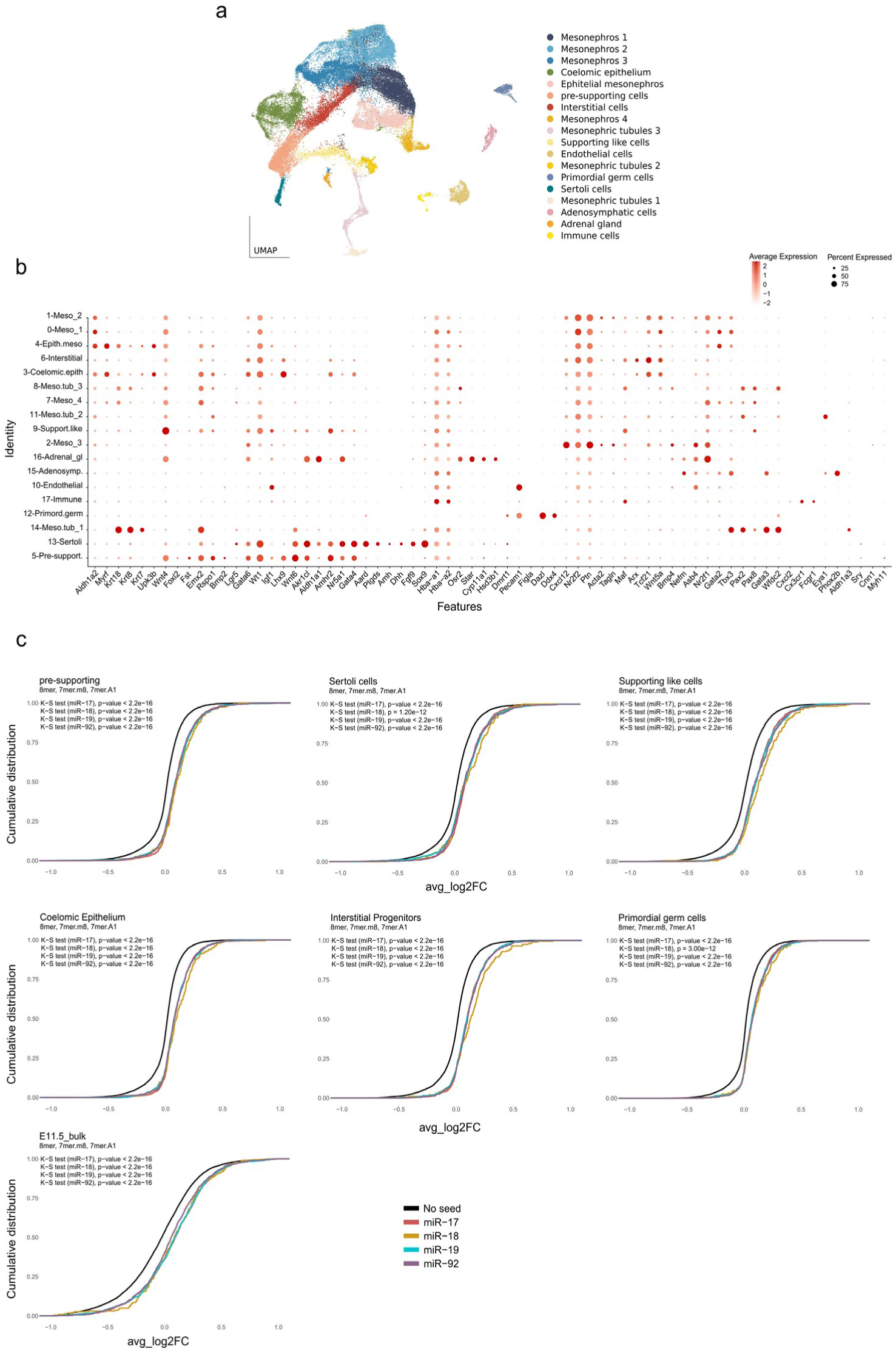
scRNA-seq clustering, annotation and Cumulative Distribution Fraction (CDF) plots of gene expression. a, UMAP representation of single-cell data from XY control (E11.5), XY *miR-17-92* KO (E11.5 and E11.75) and XX (E11.5) control mouse gonads showing initial cluster identification and annotation. b, Expression of representative marker genes and cluster annotation. c, CDF plots of log2-fold changes of genes without (black) or with (coloured) 8mer, 7mer.m8, and 7mer.A1 matches (data obtained from https://www.targetscan.org) for members of the *miR-17∼92* cluster in their 3’UTR using the bulk and scRNA-seq differential expression datasets at E11.5-E11.75.

**Extended Data Figure 4.**
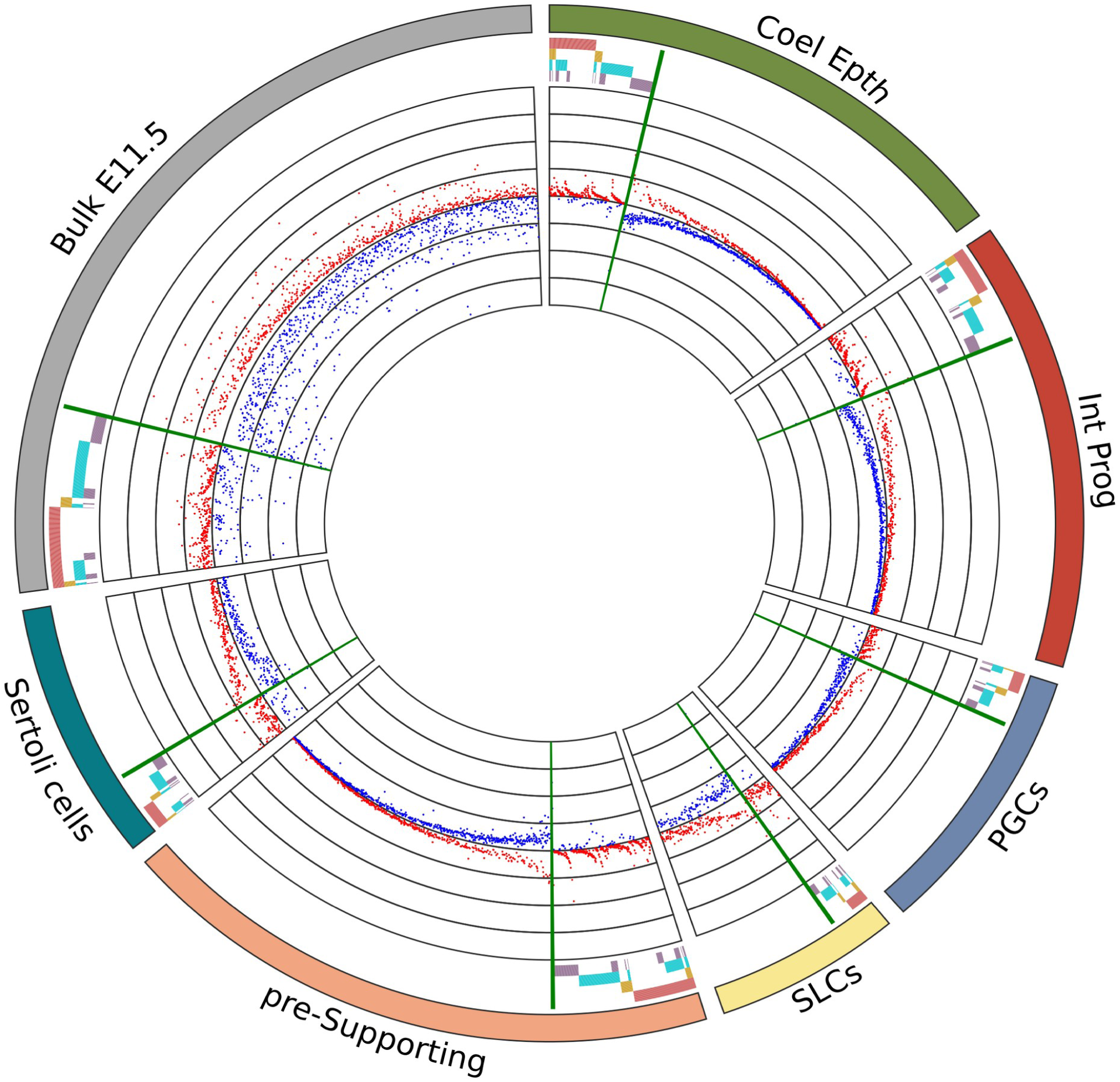
Higher resolution image of the circos plot shown in Figure 3a. Coel. Epth., Coelomic Epithelium; Int. Prog., Interstitial Progenitors; PGCs, Primordial Germ Cells; SLCs, supporting-like cells.

**Extended Data Figure 5.**
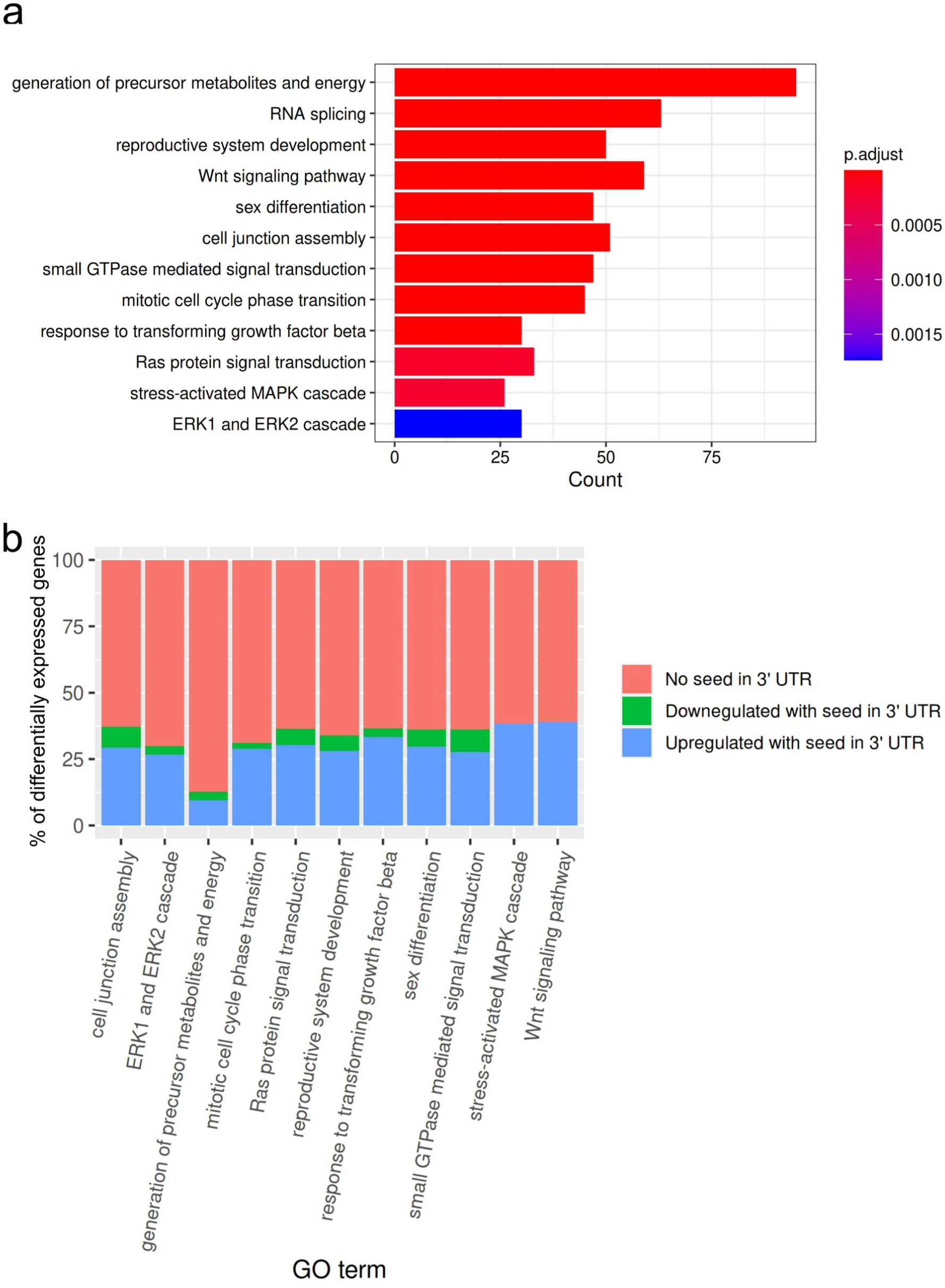
Gene Ontology analysis of mutant pre-supporting cells. **a,** GO analysis using DEG between XY control and *miR-17∼92* KO pre-supporting cells. **b,** Percentage of up-and downregulated *miR-17∼92* target genes present in the GO categories shown in **a**.

**Extended Data Figure 6.**
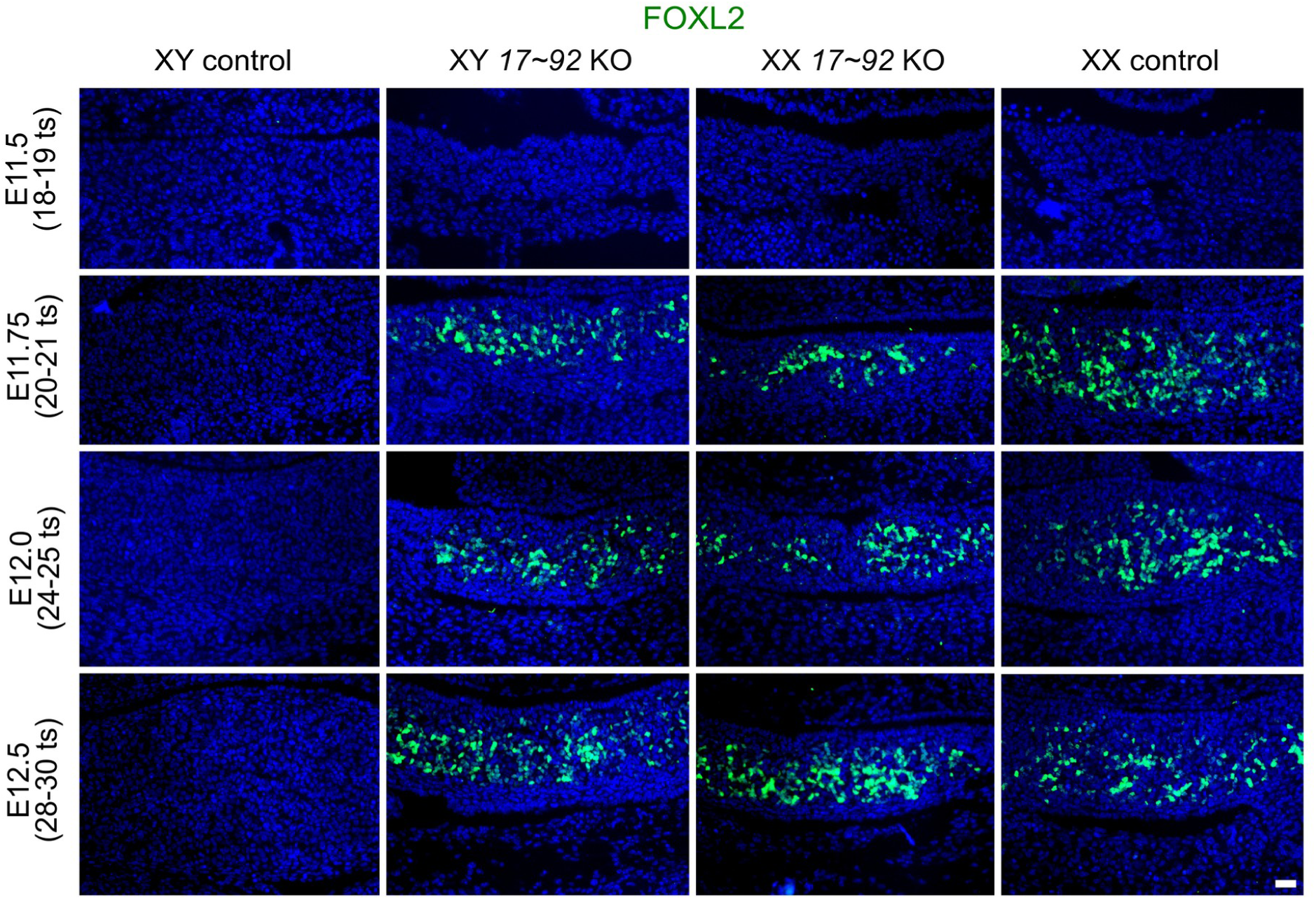
FOXL2 expression in *miR-17∼92* KO gonads. Immunofluorescence for FOXL2 in control and mutant gonads during (E11.5) and shortly after (E11.75-E12.5) the sex determination stage. Scale bar represents 50 μm.

**Extended Data Figure 7.**
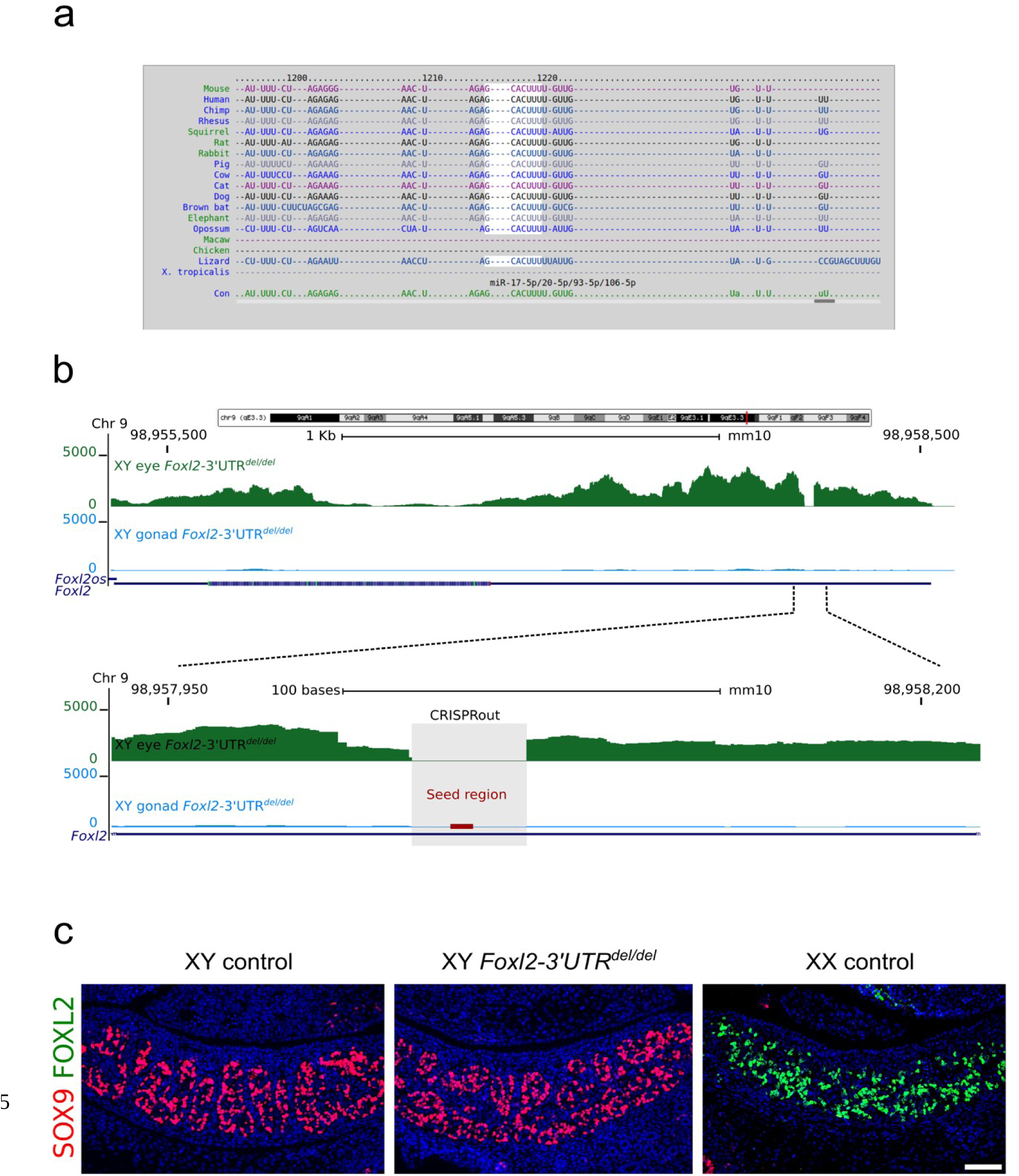
Deletion of the *miR-17* seed family binding site in the 3’ UTR of *Foxl2*. **a,** Predicted binding site for the *miR-17* seed family in the 3’ UTR of *Foxl2* using Targetscan (http://www.targetscan.org/). **b,** Transcript reads mapped to the *Foxl2* genome region from embryonic eyes (top track) and gonad (bottom track) of XY mice with a deletion of the predicted *miR-17* seed family binding site in the 3’ UTR of *Foxl2* (*Foxl2-3’-UTR^del/del^*) at E12.5 (top). Note that the 3’UTR deletion does not affect *Foxl2* transcription in tissues where the gene is expressed, like the eye. High magnification indicating the deleted region (grey rectangle) and the seed position (red bar) in the 3’ UTR (bottom). **c,** SOX9-FOXL2 double immunofluorescence in control and mutant gonads at E12.5. SOX9 is apparently normally expressed in XY *Foxl2-3’-UTR^del/del^* gonads, while FOXL2 is not present. Scale bar represents 100 μm for all figures.

**Extended Data Figure 8.**
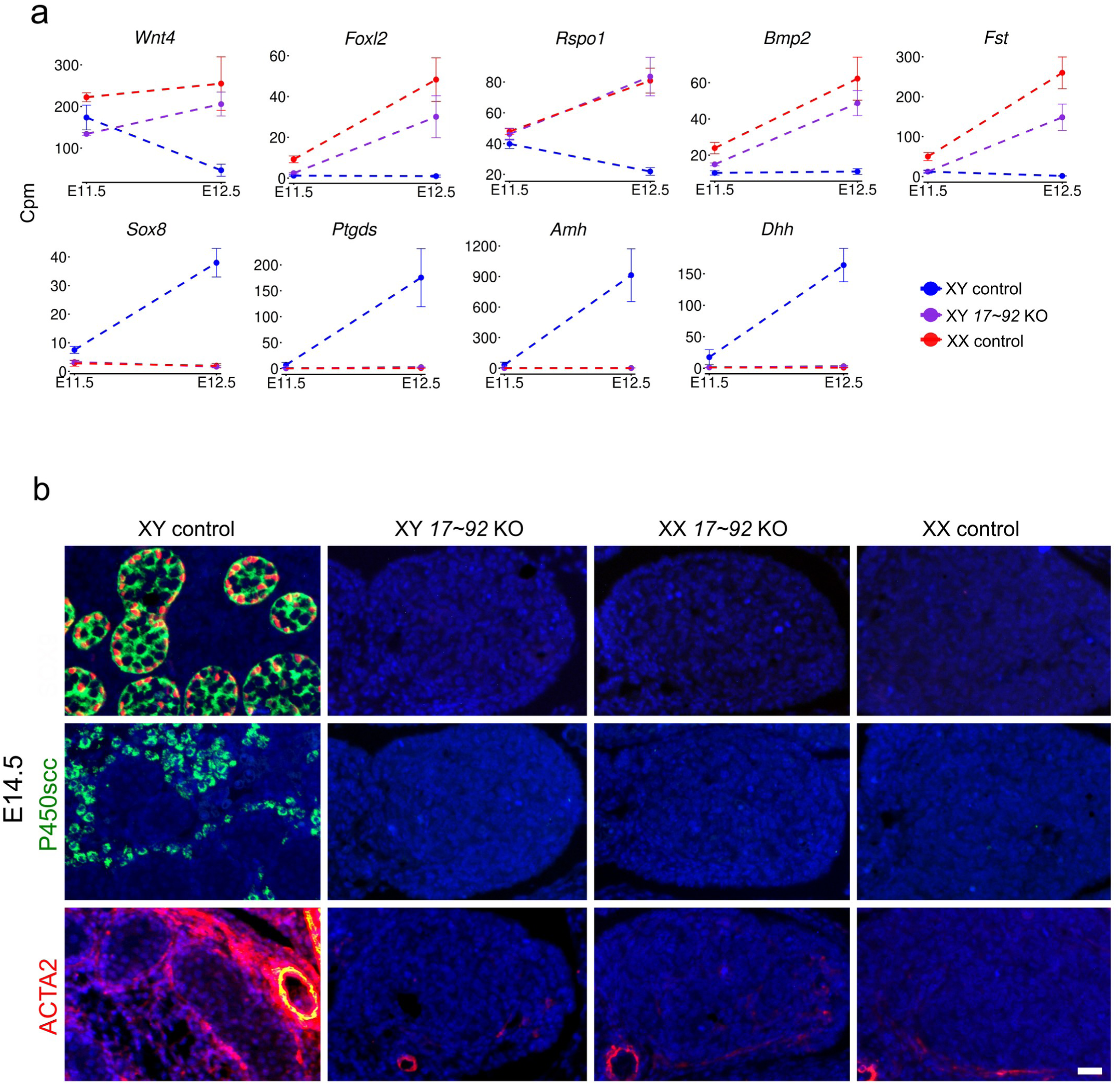
Gonadal expression of marker genes after the sex determination stage. **a,** Gonadal Bulk RNA-seq transcripts levels (in Count per million, Cpm) of ovary-specific markers (*Wnt4*, *Foxl2*, *Rspo1*, *Bmp2*, *Fst*) and testis-specific markers (*Sox8*, *Ptgds*, *Amh*, *Dhh*), in control and mutant mice at E11.5 and E12.5. **b,** Immunofluorescence for testis-specific markers in control and mutant gonads at E14.5. At this stage, XY *miR-17∼92* KO gonads show no expression for markers of Sertoli (SOX9 and AMH), Leydig (P450scc) and peritubular myoid cells (ACTA2). In addition, ACTA2 expression reveals the presence of the coelomic vessel in XY control gonads, but not in the other genotypes. Scale represents 20 μm.

**Extended Data Figure 9.**
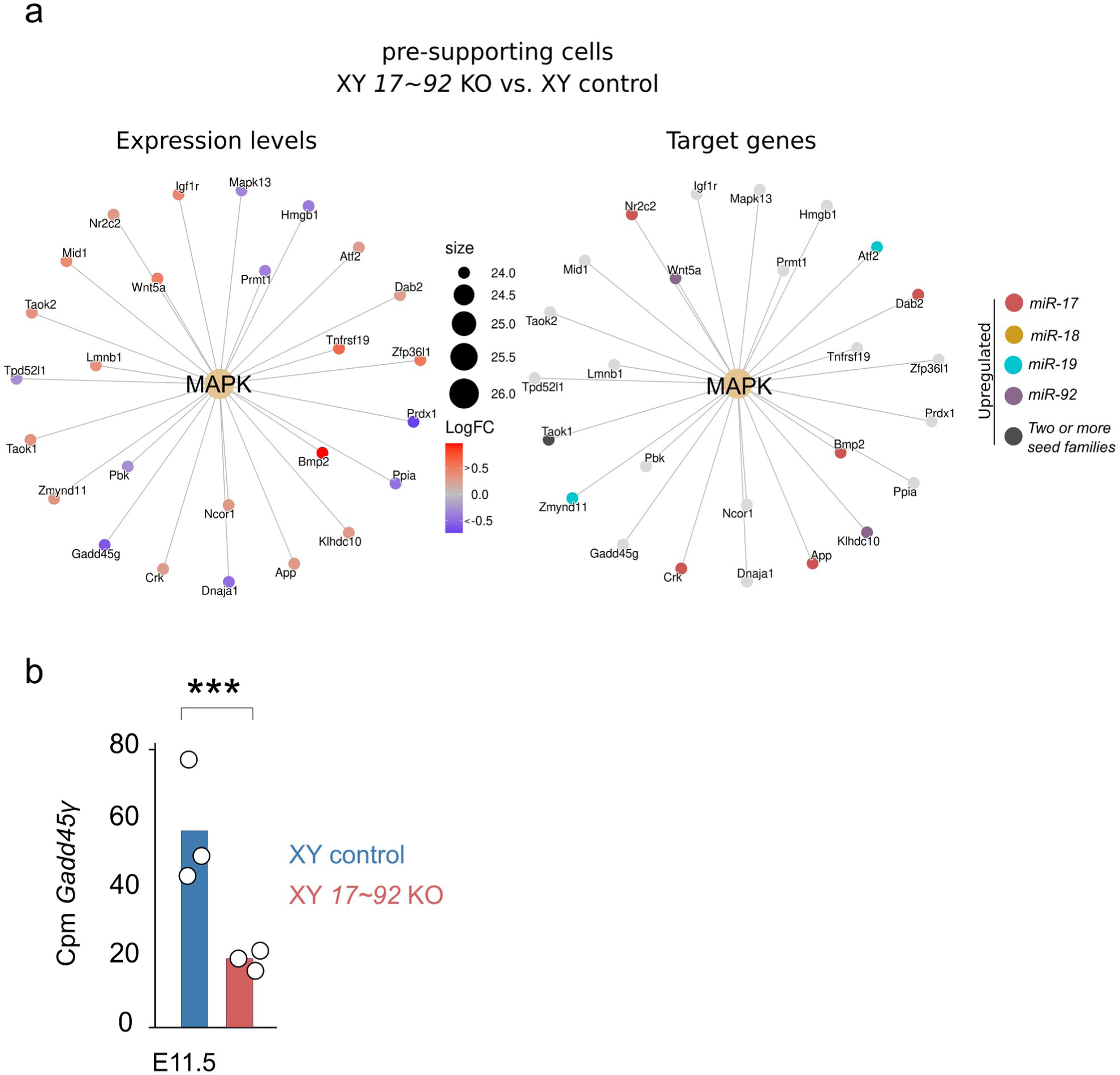
MAPK signalling pathway in the pre-supporting cells. **a**, MAPK gene-concept network using DEGs between XY mutant and control pre-supporting cells at E11.5-E11.75. Log_2_FCs (left) and predicted targets for the four *miR-17∼92* seed families (right) are depicted. **b,** Gonadal expression levels of *Gadd45γ* in control and mutant testes at E11.5 (calculated from the Cpm data of the E11.5 bulk RNA-seq dataset).

**Extended Data Figure 10.**
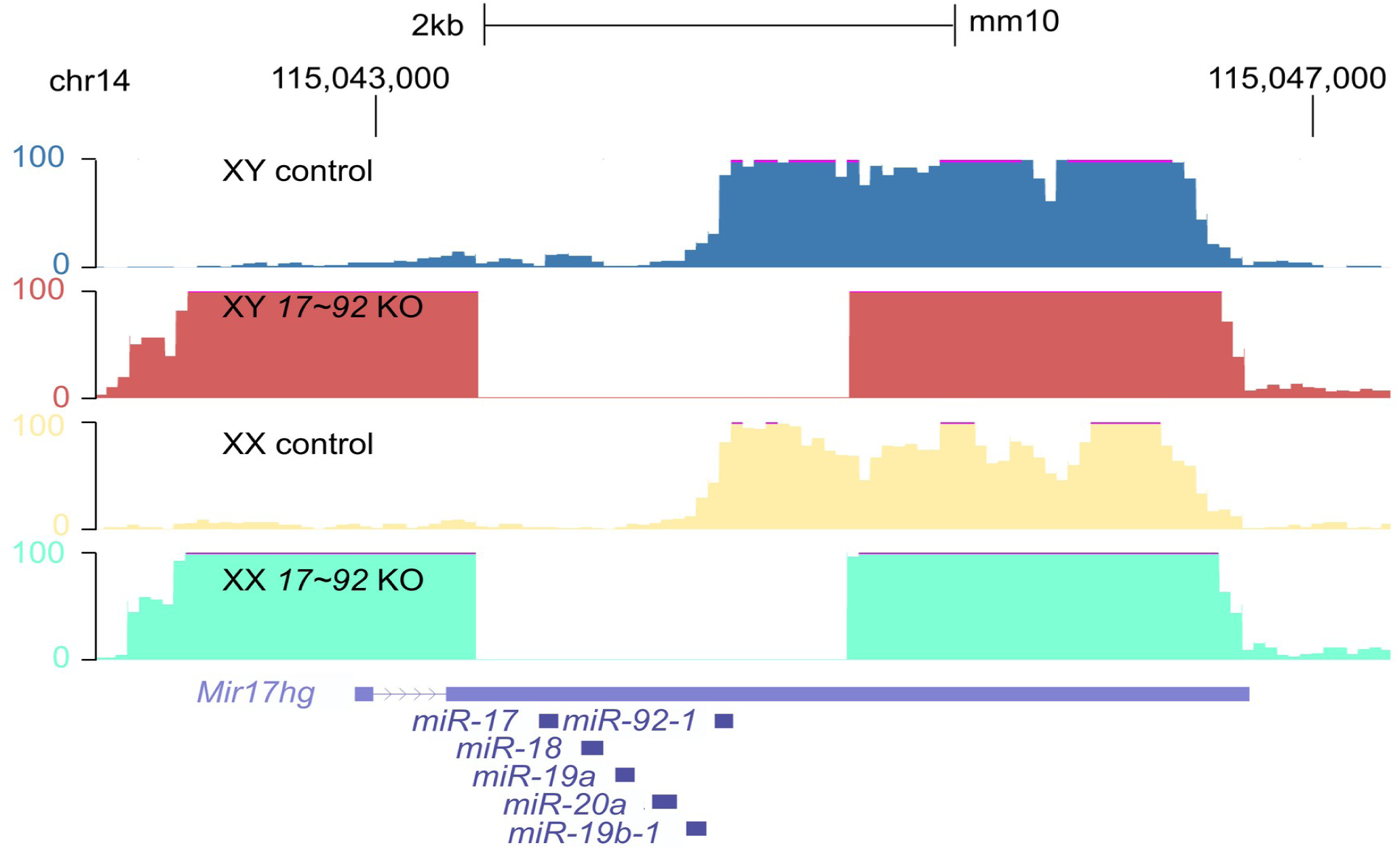
Effective deletion of the *miR-17∼92* cluster in mice. Gonadal transcript reads mapped to the *miR-17∼92* genome region of XY control (blue), XY *miR-17∼92* mutant (red), XX control (yellow) and XX *miR-17∼92* mutant (green) mice at E11.5.

